# RXR agonist S169 inhibits HBV/HDV entry *in vitro* by disrupting KIF4-dependent NTCP trafficking

**DOI:** 10.1101/2025.04.21.649881

**Authors:** Sameh A. Gad, Saied A. Hussein, Kento Fukano, Takanobu Kato, Ikuo Shoji, Kazuaki Chayama, Masamichi Muramatsu, Masanori Isogawa, Takaji Wakita, Daniel Merk, Hussein H. Aly

**Affiliations:** Department of Virology II, National Institute of Infectious Diseases; Japan Institute for Health Security; 1-23-1 Toyama; Shinjuku-ku: Tokyo 162-8640; Japan; Department of Microbiology and Immunology; Faculty of Pharmacy; Minia University; Minia 61519; Egypt; Department of Pathology; Immunology and Microbiology; Graduate School of Medicine; The University of Tokyo; Tokyo 113-0033; Japan; Department of Academic-Industrial Partnerships Promotion; Center for Clinical Sciences; Japan Institute for Health Security; Tokyo 162-8655; Japan; Division of Infectious Disease Control; Center for Infectious Diseases; Kobe University Graduate School of Medicine; 7-5-1 Kusunoki-cho; Chuo-ku Kobe 650-0017; Japan; Hiroshima Institute of Life Sciences; Hiroshima; Japan; RIKEN Center for Integrative Medical Sciences; Yokohama; Japan; Department of Infectious Disease Research; Institute of Biomedical Research and Innovation; Foundation for Biomedical Research and Innovation at Kobe; Ludwig-Maximilians-Universität München; Department of Pharmacy; Munich; Germany

## Abstract

Chronic hepatitis B (HBV) and hepatitis D (HDV) viral infections pose serious global health challenges, and the lack of curative therapies calls for the development of new antiviral approaches. Recently, we have identified the Kinesin Family Member 4 (KIF4) as a crucial regulator of the surface transport for NTCP (HBV/HDV entry receptor). Our research has shown that the pan-RXR agonist, Bexarotene, suppresses KIF4 expression and inhibits HBV/HDV infections. However, the clinical use of Bexarotene is limited by poor physicochemical and pharmacokinetic (PK) properties and significant side effects. Here we investigated chemically diverse RXR modulators with tuned activity, and identified the RXR agonist S169 as a novel HBV/HDV entry inhibitor. S169 effectively blocks HBV and HDV infections in primary human hepatocyte (PXB) cultures without evident cytotoxicity. S169’s suppressive effect on HBV infection was confirmed by reduction of secreted HBsAg and HBeAg, as well as inhibition of HBV cccDNA, pregenomic RNA, and intracellular HBV DNA. Mechanistically, S169 suppressed FOXM1-mediated KIF4 expression, leading to decreased NTCP surface levels and a marked decrease in HBV preS1 peptide binding to the hepatocyte cell surface. S169 is reported to exhibit enhanced oral bioavailability, favorable PK parameters, lower toxicity compared to Bexarotene. Interestingly, S169 induced a modest inhibition of NTCP-mediated bile acid transport up to 10 μM, which remained unchanged at higher concentrations. Furthermore, S169 significantly impeded HBV spread in HBV infected long-time cultured PHHs (PXB). We speculate that S169 can be a promising seed for the development of novel oral HBV and HDV antiviral therapeutics.

## Introduction

Hepatitis B virus (HBV) infection represents a considerable global public health issue, especially in Western Pacific and African regions, with approximately 254 million people worldwide living with chronic hepatitis B (CHB) (Ishida et al., 2015; Liang et al., 2023). CHB can progress to severe, life-threatening complications, including cirrhosis and hepatocellular carcinoma. Current treatment strategies for CHB include long-term administration of nucleos(t)ide analogs (NAs) or 1-5 years of pegylated interferon alpha (PEG-IFNα) (Pawlotsky et al., 2008), however they rarely attain functional viral cure (Feld et al., 2023; Pawlotsky et al., 2008).

HBV is a hepatotropic DNA virus belonging to the Hepadnaviridae family, noted for its strict liver tropism and distinct host specificity. The entry process of HBV into host hepatocytes involves high-affinity interaction with the specific entry receptor, sodium taurocholate co-transporting polypeptide (NTCP) on the hepatocyte cell surface (Yan et al., 2012). Hepatitis Delta virus (HDV) is a satellite RNA virus that uses HBV envelope proteins for cell entry and egress and hence follows the same entry mechanism of HBV (Sureau and Negro, 2016). The entry process is mediated by the interaction between the surface entry receptor NTCP and the myristoylated preS1 region, particularly the N-terminal 2-48 amino acids (aa), of HBV’s large surface protein (LHBs) (Barrera et al., 2005; Gripon et al., 2005; Yan et al., 2012).

NTCP-mediated HBV entry has emerged as a promising therapeutic target for developing anti-HBV drugs. Targeting this pathway offers the potential advantage of blocking HBV infection or cell-to-cell spread, thereby preventing HBV integration into the host chromosome and the establishment of HBV covalently closed circular DNA (cccDNA) minichromosome (Baumert et al., 2014; Levrero et al., 2009; Tong and Li, 2014).

Previously, we demonstrated that HBV and HDV entry can be blocked *in vitro* by targeting NTCP surface trafficking, a process primarily regulated by the motor kinesin KIF4. Although the pan-RXR agonist Bexarotene efficiently downregulates KIF4, thereby reducing NTCP surface expression and abrogating HBV infection (Gad et al., 2022), its therapeutic application is limited by poor pharmacokinetic (PK) parameters, several off-target activities, and thus undesirable side effects. In the search for improved RXR modulators, we tested a collection of chemically diverse RXR modulators (Heitel et al., 2019; Pollinger et al., 2019; Schierle et al., 2021). Among them, S169 which is a potent RXR agonist derived from the FDA-approved, non-steroidal anti-inflammatory drug (NSAID) Oxaprozin, exhibits significantly lower cytotoxicity and more favorable PK properties than Bexarotene. S169 abrogated NTCP’s surface transport and in-turn significantly inhibited HBV and HDV entry.

## Materials and Methods

A full description of the experimental procedures is available in the Supplementary Materials and Methods, while the most significant aspects are outlined briefly in this section.

### HBV and HDV infection assays

HBV (genotype D) derived from HepAD38.7-Tet cells (Gad et al., 2022) was used to inoculate primary human hepatocytes (PXB cells) at 1000 GEq/cell with 4% PEG8000. After 16 h, cells were washed and cultured for 7–15 days. HBV infection was assessed by ELISA (HBsAg, HBeAg), immunofluorescence (intracellular HBcAg), Southern blot (intracellular HBV DNA), and qPCR/RT-qPCR (cccDNA, extracellular HBV DNA, pgRNA). However, HDV used for infection was derived from Huh7 cells co-transfected with pSVLD3 and pT7HB2.7 (Gad et al., 2022). PHH (PXB) cells were inoculated at 20 GEq/cell with 5% PEG8000 for 16 h, then washed and cultured for 6 days. HDV infection was assessed by RT-qPCR quantification of HDV RNA (Gad et al., 2022).

### HBV preS1 binding assay

Attachment of TAMRA-labeled, myristoylated preS1 peptide (40 nM) to HepG2-hNTCP or PHH (PXB) cells was assessed after 30 min incubation at 37°C, followed by washing, fixation, and DAPI staining. Binding of the PreS1-TAMRA signal to the cell surface was visualized by fluorescence microscopy (Gad et al., 2022).

### HBV replication assay

HepAD38.7-Tet cells cultured without tetracycline were treated with DMSO or S169. Entecavir (10 μM) and tetracycline (0.4 μg/mL) served as positive controls. After 3 days, extracellular and intracellular HBV DNA levels were quantified by real-time PCR (Shi et al., 2023).

### HBV spread assay

PHH (PXB) cells were infected with HBV genotype C (5 GEq/cell, PhoenixBio) in the presence of 4% PEG8000 for 16 h, followed by washing to remove the unbound virus, cells were cultured for 42 days to allow viral spread. Supernatants were collected every 6 days for HBV DNA quantification (Tsukuda et al., 2017).

### Isolation and immunoblotting analysis of surface proteins

Cell surface proteins were isolated using surface biotinylation assay followed by immunoblotting analysis, as previously described (Gad et al., 2022). E-cadherin (CDH-1) was used as a loading control for surface fraction. For detection of NTCP, protein samples were treated with Peptide-N-Glycosidase F (250 U) to remove N-linked glycosylation of glycosylated NTCP prior to SDS-PAGE and immunoblot analysis (Aly et al., 2016; Gad et al., 2022).

### Cell viability assay

Cell viability assay was performed using Cell Proliferation Kit I (MTT, Roche) according to the manufacturer’s protocol.

### Data analysis and statistical evaluation

Data are presented as mean ± SD from two to three independent experiments, each performed in triplicates or quadruplicates. Statistical significance was assessed using two-tailed unpaired Student’s *t*-test (**, *P* < 0.01; ***, *P* < 0.001; NS, not significant).

## Results

### Screening for the anti-HBV activity of Oxaprozin derived RXR agonists

We previously showed that the RXR agonist, Bexarotene, suppressed HBV infection in human hepatocyte cell culture (Gad et al., 2022). Bexarotene clinical use is frequently associated with severe adverse outcomes which possibly occur due to its off-target cellular effects (Assaf et al., 2006; Lewandowski et al., 2025; Takamura et al., 2022). Aiming to overcome these limitations, we screened the anti-HBV activity of a chemically diverse set of RXR agonists; S169, JP147, JP175, and PH299 (Fig. S1) using the most physiologically relevant HBV infection model, primary human hepatocyte (PXB) cells. This selection of chemical tools covers a broad range of RXR activation including full and partial agonism as well as selective modulation. We used HBV (genotype D) infection inoculum derived from the culture supernatant of HepAD38.7-Tet cells. PHHs (PXB) were treated with the indicated compounds at 5 μM concentration 72 h prior to HBV inoculation. To investigate the effect of these compounds on HBV infection, the detection of the secreted HBsAg and the intracellular HBcAg was performed at 8 and 16 days post-infection (dpi) respectively (Fig. 1A). To validate the experimental conditions, DMSO and Bexarotene were included as negative and positive controls, respectively. While all RXR modulators showed anti-HBV activity, S169 exhibited the highest suppressive effect on HBV infection and significantly inhibited both intracellular HBcAg (Fig. 1B) and the secreted HBsAg (*P* < 0.001) (Fig. 1C) to levels comparable to Bexarotene. No detrimental effect on the cell viability was observed with any of the compounds tested (Fig. 1D).

**FIG 1.**
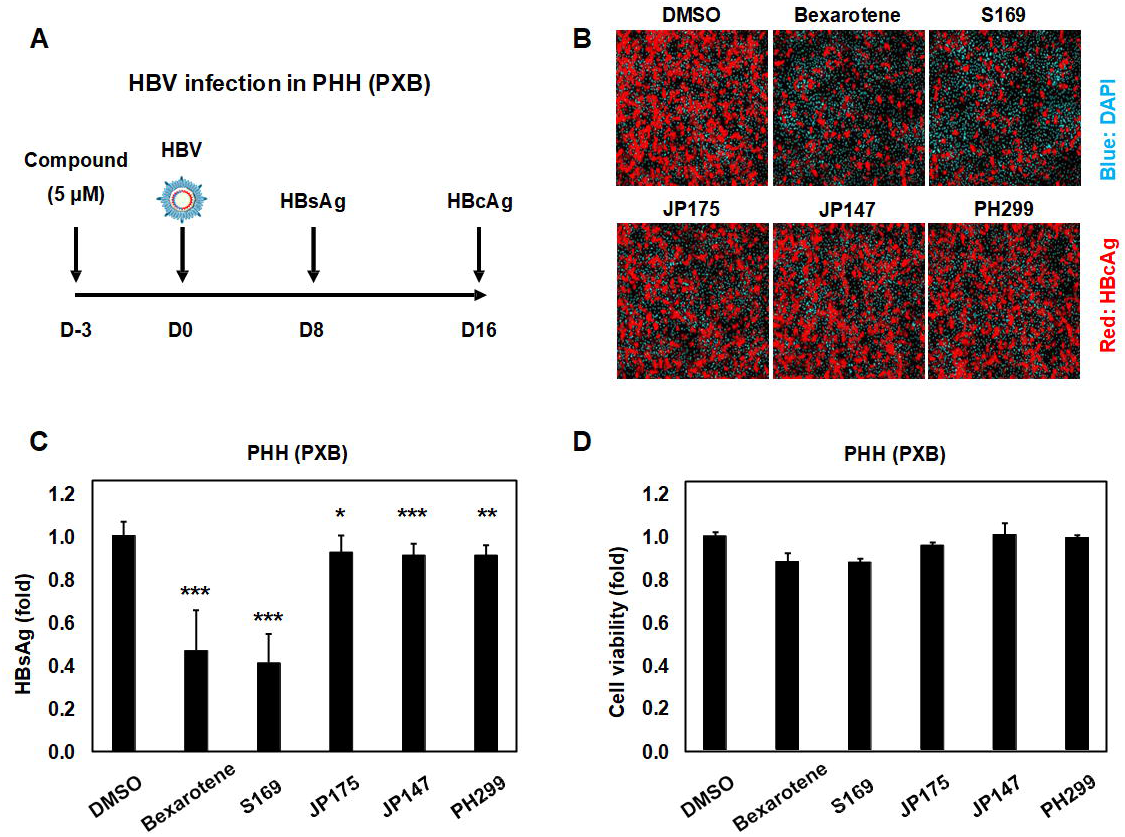
Screening for novel Oxaprozin derived RXR agonists with anti-HBV activity. **(A)** A schematic diagram showing the timeline for compound treatment and subsequent HBV infection in PHHs (PXB); PHHs were treated with the indicated compounds (5 μM) for 72 h then the compounds were withdrawn from the culture medium 3 h before HBV (genotype D) inoculation. HBV infection was performed at 1,000 GEq per cell in the presence of 4% PEG8000 for 16 h. DMSO and Bexarotene were included as negative and positive controls, respectively. After washing, the cells were cultured for an additional 15 days. **(B)** The intracellular HBcAg was detected at 16 dpi by IF. Red and blue signals show the staining of HBcAg and nucleus (DAPI), respectively. **(C)** The secreted HBsAg was quantified at 8 dpi by ELISA and the data were presented as fold changes, relative to DMSO pretreated cells. **(D)** Cell viability was evaluated at 16 dpi by the MTT assay. Data are presented as mean ± SD. *, P < 0.05; **, P < 0.01; ***, P < 0.001; NS, not significant.

### S169 efficiently blocked HBV infection in PHH (PXB) cells

We further evaluated the inhibitory effect of S169 treatment on HBV infection at different doses (Fig. 2A). S169 treatment significantly reduced permissiveness of PHHs (PXB) to HBV infection in a dose-dependent manner as indicated by the dose-dependent suppression of the secreted HBsAg (*P* < 0.001) (Fig. 2B), secreted HBeAg (*P* < 0.001) (Fig. 2C), and intracellular HBV pgRNA (*P* < 0.001) (Fig. 2D) levels; the 50% inhibitory concentration (IC_50_) was estimated to be 2.74 ± 0.8 μM, 2.05 ± 0.4 μM, and 0.99 ± 0.3 μM based on HBsAg, HBeAg, and HBV pgRNA, respectively. For validation of experimental data, DMSO and Myr-B were included as negative and positive controls, respectively. Surprisingly, PHH cultures tolerate S169 treatment over a wide range of doses, with the 50% cytotoxic concentration (CC_50_) being above 100 μM (Fig. 2E). Most importantly, S169 treatment resulted in a substantial decrease of HBV cccDNA (Fig. 2F) (*P* < 0.01) and inhibited the total intracellular HBV DNA (Fig. 2G) to levels equivalent to Myr-B treatment.

**FIG 2.**
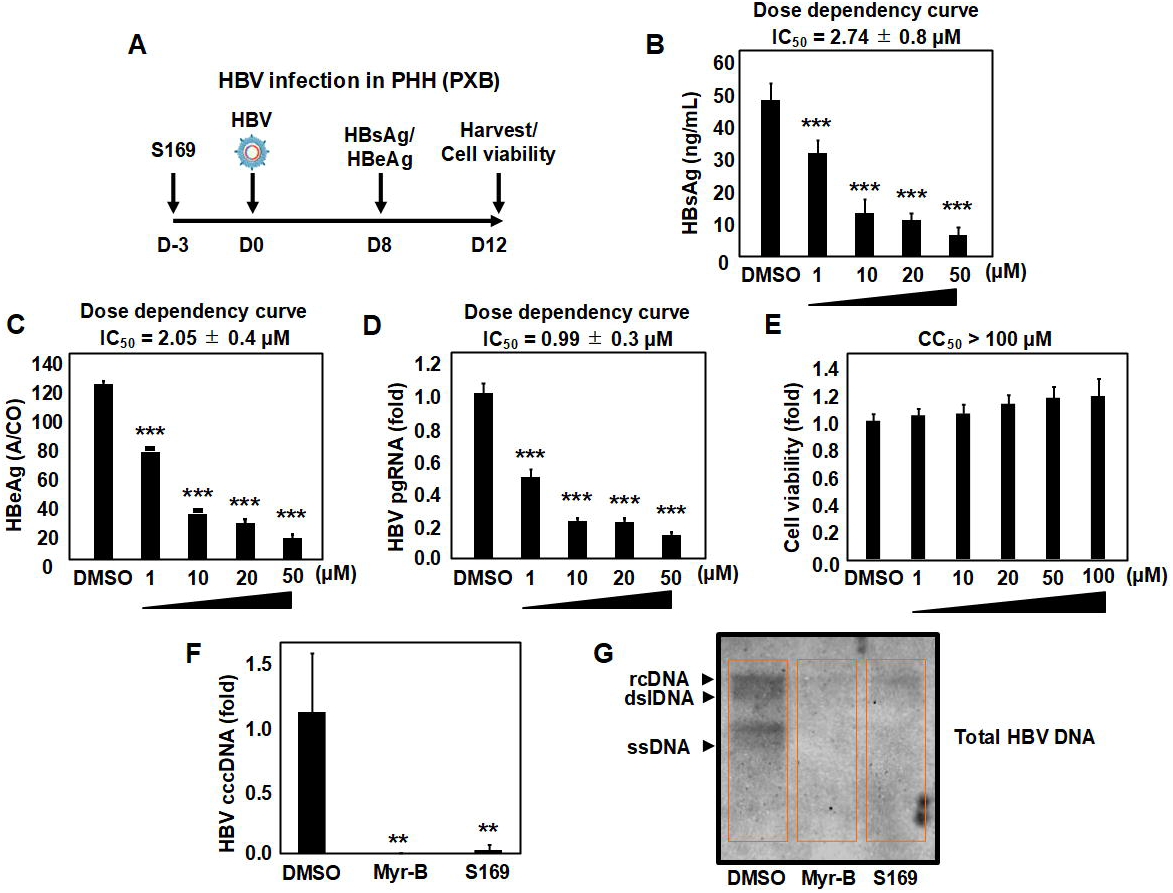
S169 efficiently suppressed HBV infection in PHH (PXB) cells. **(A)** The timeline scheme for S169 treatment and subsequent HBV infection in PHHs (PXB); The cells were treated with DMSO or different doses of S169 (1 μM, 10 μM, 20 μM, and 50 μM) for 72 h followed by withdrawal of DMSO and S169 from the culture medium 3 h before HBV inoculation (genotype D), HBV infection was conducted at inoculum (1,000 GEq/cell) in the presence of 4% PEG8000 for 16 h. DMSO pretreated cells were concomitantly treated with or without 100 nM Myr-B during HBV inoculation. After washing, the cells were cultured for an additional 11 days. **(B, C)** The secreted HBsAg and HBeAg were collected at 8 dpi, quantified by ELISA, and presented relative to the DMSO treated cells. **(D)** HBV pgRNA and **(E)** Cell viability were evaluated at 12 dpi by RT-qPCR, and MTT assay, respectively. The cells were treated with DMSO or S169 (50 μM) following the same infection scheme mentioned in **(A)** then HBV cccDNA **(F)** and total HBV DNA **(G)** were extracted at 12 dpi and analyzed by qPCR and Southern blot analysis, respectively. Data are presented as mean ± SD. **, P < 0.01; ***, P < 0.001.

### S169 hampered the early phases of HBV infection cycle

Given that S169 abrogates HBV infectivity in human hepatocytes, additional analyses were conducted to ascertain whether it selectively impacts the early stages (from viral entry to transcription) or extends to the late phases (from replication to particle release) of the HBV infection cycle. PHHs (PXB) were treated with S169 (5 μM) as follows: pre = 3 days before HBV inoculation; early post = from 1 to 5 dpi; late post = from 5 to 8 dpi; and whole = 3 days before HBV inoculation and continued from 1 to 8 dpi (Fig. 3A). The anti-HBV activity was determined by the quantification of the secreted HBeAg. Bexarotene and the entry blocker Myr-B were employed as negative and positive controls, respectively. The primary suppressive effect of S169 on HBeAg levels was observed in pre-, or whole-treatment settings whereas a modest inhibition was noted when the compound was applied in early post- or late post-treatment settings (Fig. 3B). Moreover, we utilized the HBV stably expressing cell line, HepAD38.7-Tet, which supports inducible HBV replication following tetracycline (TET) withdrawal. The nucleoside analog entecavir (ETV), which inhibits HBV replication, served as a positive control. We found that S169 treatment at various concentrations did not affect intracellular or extracellular HBV DNA levels (Fig. 3C), indicating that S169 does not target viral replication (Kang et al., 2019; Shi et al., 2023). In addition, we tested the compound effect on the HBV cccDNA synthesis through the intracellular recycling pathway of viral nucleocapsids in HepAD38.7-Tet cell line (Tang et al., 2019). Following the withdrawal of TET, treatment with S169 at concentrations of 5 μM and 20 μM for 6 days did not exhibit a significant repressive effect on HBV cccDNA levels, as measured by qPCR (Fig. 3D). These findings strongly suggest that S169 exerts its effects during the initial stages of HBV infection, likely preceding nuclear import and establishment of nuclear HBV cccDNA.

**FIG 3.**
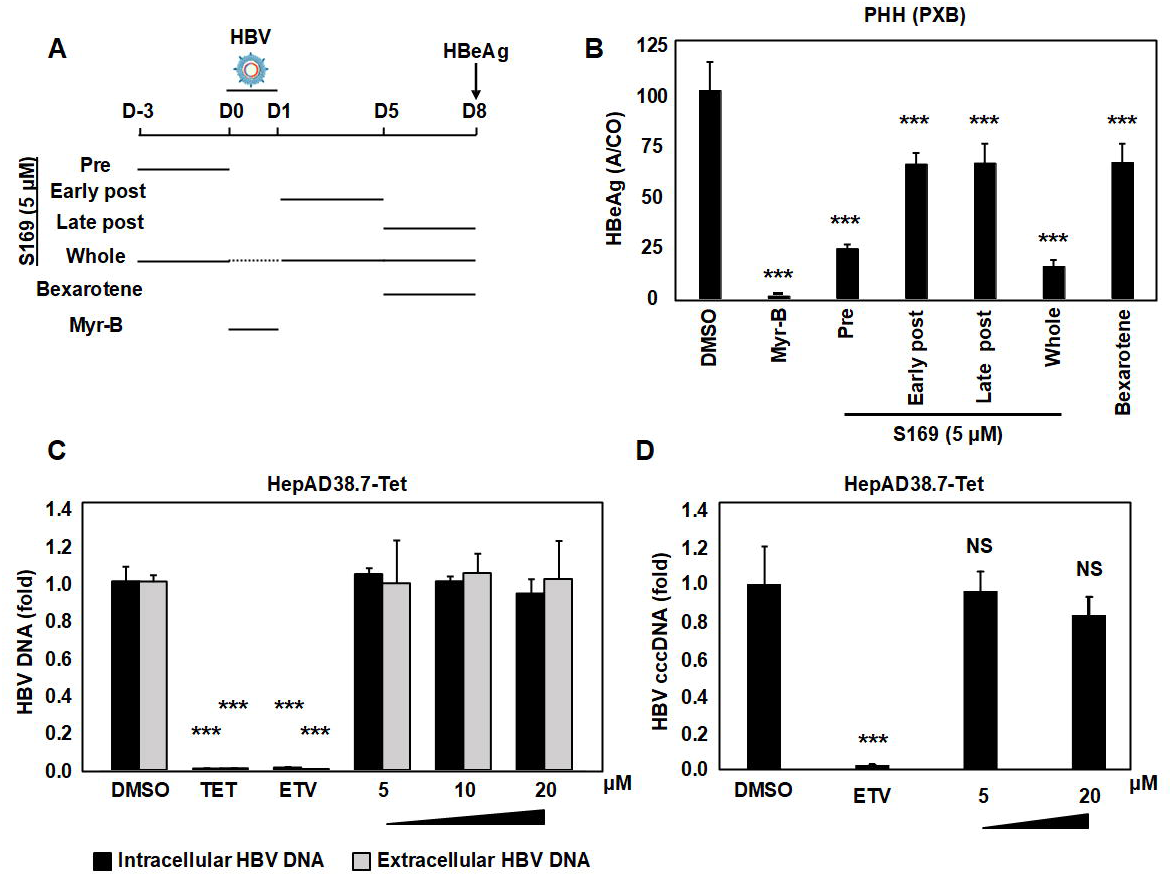
S169 restricts the early phases of HBV infection cycle. **(A)** A schematic diagram showing HBV infection protocol; PHHs (PXB) were treated with 5 μM S169 at different time schedule (pre, early post, late post, and whole) as shown in the figure. HBV (genotype D) inoculation was performed at 1,000 GEq per cell in the presence of 4% PEG8000 for 16 h. DMSO pretreated cells were concomitantly treated with or without Myr-B (100 nM) during HBV inoculation. Bexarotene (another RXR agonist) treated group was also included in which Bexarotene was added at 5 μM (from 5 to 8 dpi). **(B)** The secreted HBeAg were harvested at 8 dpi, evaluated by ELISA, and the data were presented as absorbance/cut-off (A/CO) values relative to the DMSO treated cells. **(C)** HBV-pgRNA transcription was initiated in HepAD38.7-Tet cells by stopping tetracycline treatment. DMSO or S169 were added at the indicated concentrations. Entecavir (ETV, 10 µM) and Tetracycline (TET, 0.4 μg/mL) were also used to suppress HBV-DNA reverse transcription, or HBV-pgRNA transcription respectively. At 3 days post-treatment, both total intracellular HBV DNA (*black bars*) and extracellular HBV DNA (*gray bars*) released in the culture supernatant were evaluated by real time PCR. **(D)** Transcription of HBV-pgRNA was initiated in HepAD38.7-Tet cells by tetracycline withdrawal, and the cells were treated with DMSO, S169 or Entecavir at the indicated concentrations. At 6 days post-treatment, HBV cccDNA was extracted in proteinase K free conditions and quantified by real time PCR. The data were pooled to assess the statistical significance and presented as mean ± SD. ***, *P* < 0.001; NS, not significant.

### S169 suppressed HBV and HDV entry into PHH (PXB) cells

The binding of the preS1 region of HBV’s large surface protein (LHBs) to the hepatocyte specific HBV entry receptor, NTCP (Yan et al., 2012), is essential for HBV entry and infection. To further analyze the role of S169 in this early stage, we examined the impact of S169 treatment on this specific binding using HBV preS1 binding assay where the binding efficiency of a fluorescence-labeled preS1 peptide (preS1-TAMRA) to S169 pretreated HepG2-hNTCP cells were detected by IF (Fig. 4A). Myr-B, which competitively inhibits preS1-TAMRA binding to NTCP, was employed as positive control. S169 treatment for 72 h dramatically suppressed the interaction between preS1-probe and surface NTCP (as shown in Fig. 4A). Moreover, PHH (PXB) cells were used to confirm the inhibitory effect of S169 on HBV preS1 binding to the surface NTCP receptor in a more physiologically relevant paradigm (Fig. 4B). As expected, S169 treatment abrogated the fluorescent preS1-probe signals demonstrated on the PHH (PXB) cell surface (Fig. 4B). HDV is a satellite virus that uses HBV envelope proteins for cell surface NTCP interaction and hepatocyte entry, and hence follows the same entry mechanism of HBV (Sureau and Negro, 2016). S169 pretreatment suppressed PHH (PXB) permissiveness to HDV infection as indicated by significant inhibition of HDV RNA (*P* < 0.001) measured by RT-qPCR (Fig. 4C). These findings suggest that S169 inhibited the NTCP-mediated entry of HBV and HDV.

**FIG 4.**
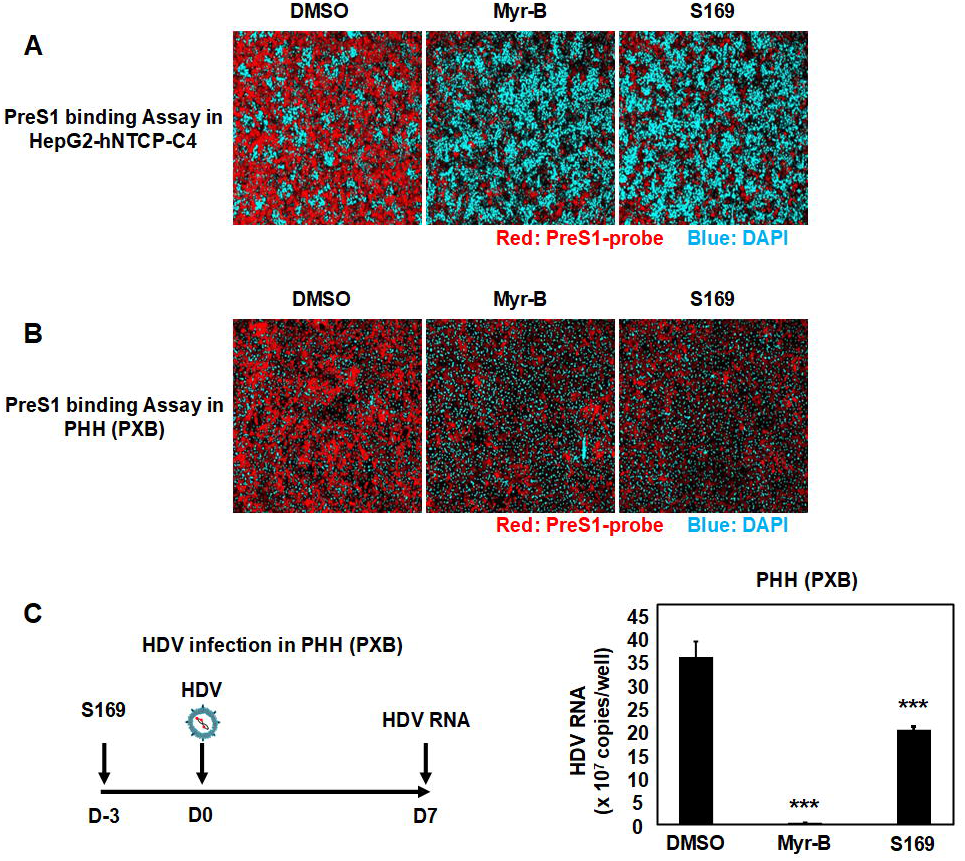
S169 suppressed HBV and HDV entry into human hepatocytes. **(A)** HepG2-hNTCP cells were treated with DMSO or S169 (20 µM) for 72 h and then incubated with 40 nM TAMRA labeled preS1 peptide (preS1 probe) for 30 min at 37°C. DMSO treated cells were incubated with the preS1 probe either in the absence or presence of Myr-B (100 nM) **(B)** PHHs (PXB) treated with DMSO or S169 (50 µM) were subjected to HBV preS1 binding assay (as described above). Red and blue signals indicate the preS1 probe and the nucleus, respectively in both **(A)** and **(B)**. **(C)** The left-sided diagram depicts the HDV infection scheme in PHHs (PXB); PHHs treated with DMSO or S169 (50 µM) for 72 h were inoculated with HDV at 20 GEq/cell 3 h following DMSO and S169 withdrawal from the culture medium. DMSO pretreated cells were concomitantly treated with or without 100 nM Myr-B to compete with HDV binding to NTCP during the viral inoculation. HDV RNA level was extracted from the cells at 7 dpi, quantified by RT-qPCR and presented relative to the DMSO treated cells (right panel). The data were presented as mean ± SD. ***, *P* < 0.001.

### S169 inhibited the KIF4-mediated transport of NTCP to the cell surface

As we previously reported that compounds with RXR agonistic activity inhibit KIF4-mediated NTCP transport to the cell surface by downregulating the FOXM1-regulated KIF4 expression, thereby suppressing the entry of HBV and HDV into hepatocytes (Gad et al., 2022), we analyzed the impact of S169 treatment on FOXM1 and KIF4 expression levels in PHHs (PXB). Immunoblotting analysis showed that S169 treatment markedly reduced FOXM1 and KIF4 expressions (Fig. 5A). Since KIF4 is essential for cell surface transport of NTCP, we analyzed the effect of S169 on the levels of cell membrane-associated NTCP. Cellular fractionation revealed that although treatment with S169 did not affect the overall cellular expression of NTCP (Input, Fig. 5B), it profoundly reduced the level of NTCP protein in the cell surface fraction (Surface, Fig. 5B), as determined by immunoblotting. Surface NTCP is implicated in sodium-dependent intrahepatic bile acid uptake (Anwer and Stieger, 2014). Therefore, a reduction in surface NTCP levels is expected to impair the uptake of bile acids into hepatocytes. Accordingly, we observed that S169 treatment at various doses significantly suppressed Na+-dependent bile salt uptake (*P* < 0.001). The S169 suppression of Na+-dependent bile salt uptake showed a plateau for doses beyond 10 μM (Fig. 5C). It is also worth noting that the suppression mediated by S169 on Na+-dependent bile salt uptake was much lower than the effect of the HBV entry inhibitor Myr-B. Overall, our data indicate that S169 targets NTCP surface transport by suppressing the expressions of the transactivator FOXM1 and its downstream KIF4 leading to significant suppression of NTCP-mediated HBV entry with a relatively lower impact on Na+-dependent bile salt uptake.

**FIG 5.**
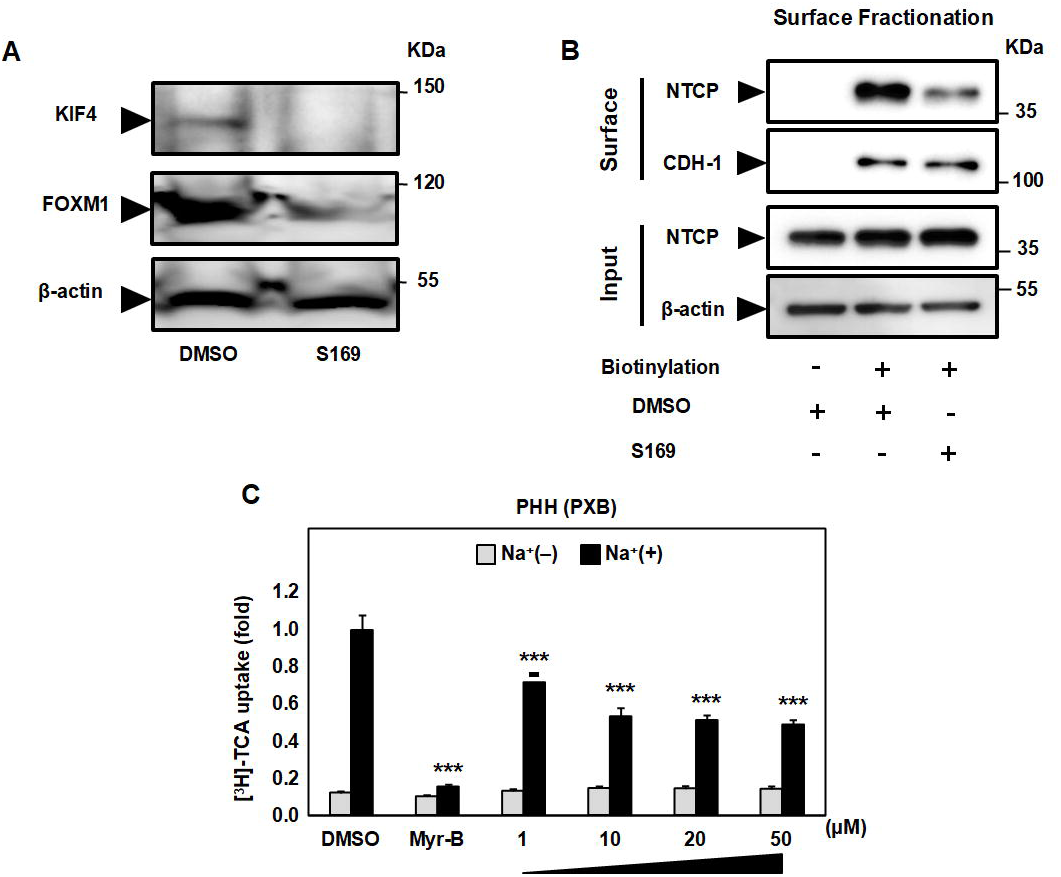
S169 suppressed KIF4-mediated NTCP surface transport. **(A)** PHHs (PXB) treated with DMSO or S169 (50 μM) for 72 h were lysed, and total cell lysate was subjected to immunoblotting to detect the expression levels of KIF4 (*upper panel*), FOXM1 (*middle panel*), and β-actin (loading control) (*lower panel*) with their corresponding antibodies. **(B)** HepG2-hNTCP cells treated with DMSO or S169 (20 μM) for 72 h, were surface biotinylated or PBS treated prior to cell lysis, then an aliquot (1/10 volume) of the cell lysate was used for Western blotting detection of total NTCP protein (*Input; upper panel)* and β-actin (loading control) (*Input; lower panel*). The remaining lysate (9/10 of the original volume) was incubated with pre-washed SA beads to capture the biotinylated surface proteins; after SA beads washing and protein elution, the surface fraction was subjected to immunoblotting to verify surface NTCP (*Surface*; *upper* panel) and CDH-1 (loading control for surface fraction) (*Surface*; *lower panel*) protein levels. **(C)** PHHs (PXB) pretreated with DMSO or different doses of S169 for 72 h were incubated with [^3^H]-taurocholic acid (TCA) at 37°C for 15 min. After washing and cell lysis, the intracellular radioactivity was measured and plotted as fold changes, relative to control DMSO treated cells to assess both sodium ion (Na^+^)-dependent (*black bars*) and Na^+^-free (*gray bars*) [^3^H]-TCA uptake. Na^+^-dependent and Na^+^-free [^3^H]-TCA uptakes were evaluated in DMSO treated cells either in the absence (negative control) or presence (positive control) of Myr-B (100 nM). The data were presented as mean ± SD. ***, *P* < 0.001.

### S169 impeded the cell-to-cell spread of HBV in long-term PHH (PXB) cultures

PHHs (PXB) have previously been shown to support HBV spread from infected hepatocytes to naïve populations (Tsukuda et al., 2017). Being an HBV entry inhibitor, we evaluated the ability of S169 to mitigate HBV (genotype C) spread in PHHs (PXB) inoculated with a low HBV inoculum (5 GEq/cell) and maintained in culture for 42 dpi to allow viral dissemination over time. S169 treatment was initiated at 6 dpi to ensure that HBV entry and initial infection had been established prior to drug administration (Fig. 6A). To monitor HBV spread, we quantified HBV DNA levels in the culture supernatant over time using qPCR. In the untreated group, extracellular HBV DNA levels increased sharply over time, whereas in the S169-treated group, HBV DNA levels remained significantly lower and showed minimal increase (Fig. 6B). Furthermore, immunofluorescence staining for HBcAg at 42 dpi revealed a substantial reduction in HBcAg signal in the S169-treated group compared to the untreated group (Fig. 6C), indicating that S169 effectively inhibits viral spread from infected cells to naïve populations.

**FIG 6.**
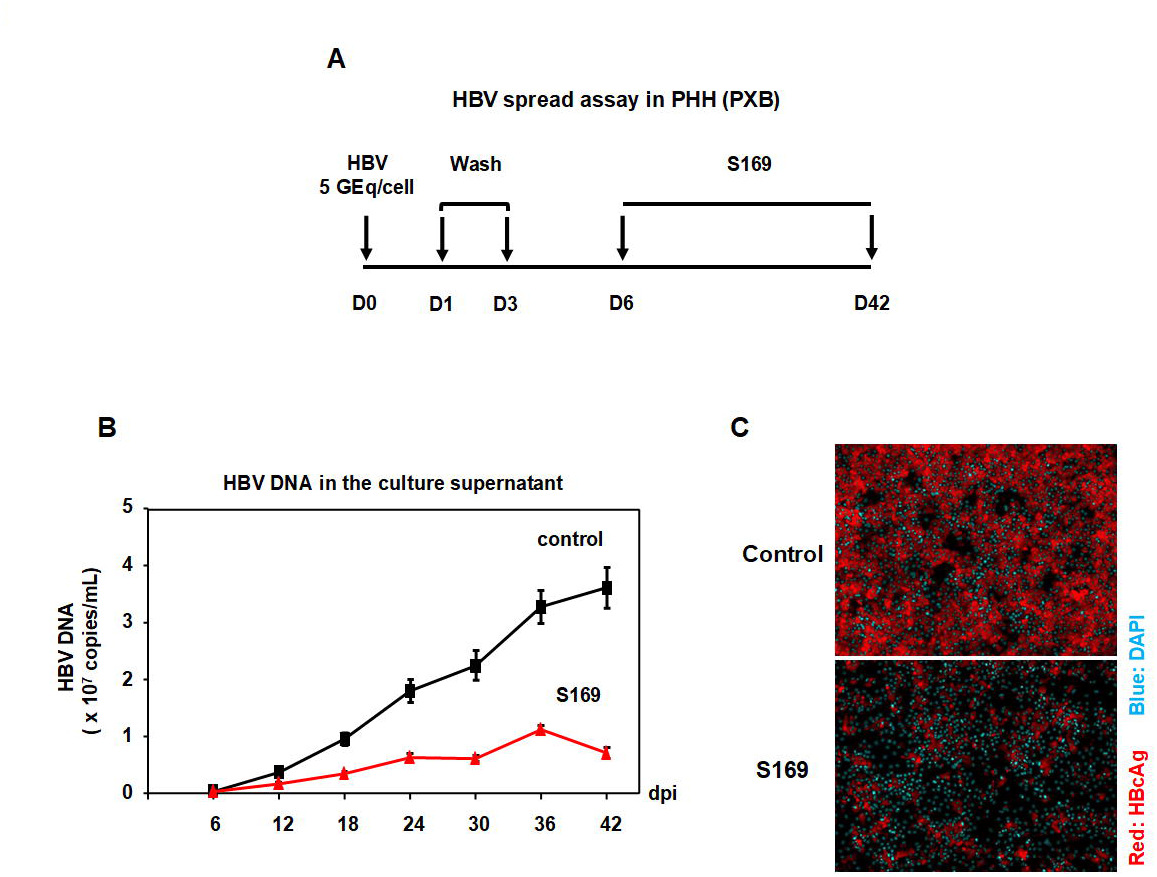
S169 blocked HBV spread in long-term PHH (PXB) cultures. **(A)** A schematic presentation showing the timeline for HBV spread assay in PHHs (PXB); PHHs were inoculated with HBV (genotype C) at very low inoculum (5 GEq/cell) in the presence of 4% PEG8000 for 16 h. DMSO or S169 (5 μM) were continuously added from 6 dpi and kept in culture for 42 days. The culture supernatant was collected every 6 days to detect extracellular HBV DNA **(B)** over time. **(C)** HBcAg was detected at 42 dpi by Immunofluorescence. Red and blue signals depict the staining of HBcAg and nucleus (DAPI), respectively.

## Discussion

Current therapeutic regimens for hepatitis B still fail to achieve a complete cure of the viral infection due to their limited impact on cccDNA pool (Yardeni et al., 2023). New strategies are needed to improve therapeutic outcomes. HBV entry inhibition has gained major interest over the last few years as an appealing approach for limiting the viral spread (Ishida et al., 2015; Tsukuda et al., 2017; Volz et al., 2013), and preventing *de novo* HBV infection in certain clinical settings. These include preventing HBV recurrence following liver transplantation, blocking vertical transmission, and enabling postexposure prophylaxis (Tu and Urban, 2018; Watashi et al., 2014).

We recently reported that the RXR agonist Bexarotene inhibits HBV/HDV entry by downregulating KIF4 expression, thereby disrupting KIF4-mediated surface transport of NTCP (Gad et al., 2022). Although Bexarotene is approved for cancer therapy (Breneman et al., 2002; Heald et al., 2003), its clinical use is confined to second- or third-line treatment due to its exceptional lipophilicity, inadequate PK and physicochemical properties, broad off-target activity, and associated severe adverse effects—including hypertriglyceridemia, hypercholesterolemia, xeroderma, and hypothyroidism (Assaf et al., 2006; Farol and Hymes, 2004). These drawbacks significantly diminish its suitability as a therapeutic candidate for HBV entry inhibition. Consequently, there is a pressing need to develop novel RXR modulators with improved pharmacological profile and lower cytotoxicity (Heitel et al., 2019; Pollinger et al., 2019; Schierle et al., 2021).

Our *in vitro* evaluation of candidate RXR ligands for anti-HBV efficacy and safety identified the RXR agonist S169 as a promising antiviral compound (Fig. 1). While S169 is structurally distinct from Bexarotene, both compounds exhibit comparable potency across RXR subtypes and nearly identical anti-HBV activity (Bexarotene IC₅₀: 1.89 μM; S169 IC₅₀: 2.74 μM for HBsAg) (Gad et al., 2022; Schierle et al., 2021), highlighting RXR activation as the common mechanism driving their antiviral effects. Notably, S169 demonstrates high selectivity for RXRs, showing no activation of RARα, PPARs, LXRs, FXR, or CAR, nor any antagonistic effect on their intrinsic activity. Moreover, in contrast to Bexarotene, S169 exhibits markedly improved oral bioavailability, superior PK and physicochemical properties, and enhanced metabolic stability. Moreover, a preliminary PK study by Schierle *et al*. following single oral administration (10 mg/kg) in mice confirmed S169’s favorable profile, with a half-life exceeding 3 hours and no evidence of hepatic accumulation, supporting S169 as a compelling lead compound for the development of HBV/HDV entry inhibitors.

HBV entry inhibitors possess the primary advantage of exerting pan-genotypic inhibition of HBV infection (Shimura et al., 2017). Accordingly, S169 was found to suppress HBV GTD infections (Fig. 2A-G; Fig. 3A and 3B) and HBV GTC viral spread (Fig. 6) in PHH (PXB) cultures. Although the main inhibitory effect of S169 and Bexarotene was on viral entry, the slight reduction in HBeAg levels observed with S169 and Bexarotene (in post-treatment settings) (Fig. 3A and 3B) can be explained by the weak inhibitory effect of RXR agonists on viral transcription from HBV cccDNA (Nkongolo et al., 2019).

HBV LHBs is proposed to form multimeric complexes, such as trimers, with NTCP. Functional inactivation of these complexes can be achieved by blocking a single NTCP subunit using direct preS1-binding entry inhibitors, indicating that full NTCP occupancy is not required for HBV entry inhibition (Li and Urban, 2016; Schulze et al., 2010). Consequently, HBV entry can be suppressed at lower inhibitor concentrations than those needed to interfere with NTCP-mediated taurocholate (TCA) transport (Nkongolo et al., 2014; Urban et al., 2014). S169 reduces NTCP surface availability, thereby limiting the multivalent interaction with LHBs and preventing productive HBV entry (IC₅₀: 2.74 μM for HBsAg; 2.05 μM for HBeAg), without significantly impairing TCA uptake (IC₅₀ ≥ 20 μM). Interestingly, TCA transport inhibition by S169 remains modest up to 10 μM (about 50 % inhibition), beyond which a plateau is observed, indicating a lack of further dose-dependent inhibition (Fig. 5C). This plateau effect may likely be attributed to the presence of KIF4-independent NTCP trafficking pathways (Anwer et al., 2005; Webster et al., 2002) that sustain a residual level of surface NTCP, even at higher S169 concentrations. In contrast, direct NTCP–preS1 interaction blockers exhibit continuous, dose-dependent suppression of TCA uptake (Ni et al., 2014; Watashi et al., 2014). These findings underscore a favorable therapeutic window for S169, enabling selective HBV entry inhibition while largely preserving physiological NTCP function.

While small molecules targeting transcriptional regulators may influence multiple downstream pathways and potentially elicit off-target effects, it is notable that, unlike the pronounced cytotoxicity reported for bexarotene at relatively low concentrations (Schierle et al., 2021), S169 exhibited no detectable impact on cell viability at doses up to 100 μM in this study. Nonetheless, future *in vivo* investigations are warranted to fully assess its safety profile and exclude potential adverse effects. Functional cure of chronic hepatitis B (CHB), indicated by HBsAg loss after six months off therapy, is infrequently attained with current therapeutic regimens (Wong et al., 2022). Combination therapies with anti-viral compounds or immune modulators which can eliminate infected hepatocytes while preventing new infections and viral spread using efficient entry inhibitors like S169 are expected to help the establishment of functional cure in patients with CHB (Lim et al., 2023).

## Aknowledgement

We gratefully acknowledge Dr. Stephan Urban at University Hospital Heidelberg for providing Myrcludex-B; and Dr. John Taylor at the Fox Chase Cancer Center, the USA for providing pSVLD3 plasmid. This study was supported by research funding from the Research Program on Hepatitis from the Japan Agency for Medical Research and Development, AMED (grants number: 22fk0310513h0001 to Kazuaki Chayama; and 24fk0310513j0703; 24fk0310503j0103, 24fk0310507j0903 to Hussein H. Aly), and from the Ministry of Education, Culture, Sports, Science and Technology, Grants in aid for Scientific Research C, 23K06575 to Hussein H. Aly.

