## Supplementary Materials for "RXR agonist S169 inhibits HBV/HDV entry *in vitro* by disrupting KIF4-dependent NTCP trafficking"

**Supplementary Materials and Methods**

**Cell lines**

Primary human hepatocytes (PXB cells) were purchased from PhoenixBio (Japan) (Ishida et al., 2015). HepG2-hNTCP-C4 and HepAD38.7-Tet cell lines were cultured in DMEM/F-12 GlutaMax media (Gibco) as previously described (Gad et al., 2022; Ibrahim et al., 2021). G418 (400 µg/ml) was added during the culture of HepG2-hNTCP-C4 cells for maintenance of hNTCP expression, HepAD38.7-Tet cells were treated with tetracycline (0.4 μg/ml) which was withdrawn from the culture medium upon induction of HBV replication.

**Peptides and Compounds**

PreS1 probe, a peptide spanning the 2-48 amino acids of the L-HBsAg preS1 region with myristoylation and 6-carboxytetramethylrhodamine (TAMRA) labeling at N- and C-termini, respectively, was synthesized by Scrum, Inc. Myrcludex-B was kindly provided by Dr. Stephan Urban at Heidelberg University. Bexarotene, Dimethyl Sulfoxide (DMSO), and Entecavir (ETV) were purchased Sigma-Aldrich; and the Oxaprozin derived RXR agonists; S169, JP147, JP175, and PH299 were synthesized as previously described (Heitel et al., 2019; Pollinger et al., 2019; Schierle et al., 2021). EZ-Link™ Sulfo-NHS-LC-Biotin (A39257) was purchased from Invitrogen.

**HBV infection assay**

HBV (genotype D) prepared from the culture supernatant of HepAD38.7-Tet cells as previously described (Gad et al., 2022) was used as the inoculum for HBV infection experiments. PHH (PXB) cells were inoculated with HBV at 1000 genome equivalent (GEq)/cell, in the presence of 4% PEG8000. After 16 h. the cells were washed out to remove the free viral particles and cultured for additional 7-15 days. HBV infection was evaluated by detecting secreted HBsAg and HBeAg (ELISA), intracellular HBcAg (Immunofluorescence), intracellular HBV DNA (Southern blot), HBV cccDNA and extracellular HBV DNA (real time PCR), and intracellular HBV pgRNA (real time RT-PCR).

**HDV infection assay**

HDV used in the infection assay was derived from the culture supernatant of Huh7 cells co-transfected with pSVLD3 and pT7HB2.7 as previously described (Gad et al., 2022). PHH (PXB) cells were infected with HDV at 20 GEq/cell in the presence of 5% PEG8000 for 16 h. The cells were then washed out to remove the free virus and cultured for an additional 6 days. HDV infection was assessed by quantification of HDV RNA by RT-qPCR.

**HBV preS1 binding assay**

Attachment of HBV preS1 peptide to HepG2-hNTCP or PHH (PXB) cell surface was evaluated by incubating the cells with 40 nM C-terminally TAMRA labelled and N-terminally myristoylated preS1 peptide for 30 min. at 37˚C. After washing, fixation with 4% paraformaldehyde and staining with DAPI, preS1 peptide binding to the cell surface was observed by fluorescent microscopy (KEYENCE, BZ-X710) (Gad et al., 2022).

**HBV replication assay**

In the absence of tetracycline, HepAD38.7-Tet cells were treated with DMSO or S169. Entecavir (10 µM) and Tetracycline (0.4 μg/mL) were used as positive controls. At 3 days post-treatment, the cell culture supernatant was harvested to detect extracellular HBV DNA level (Shi et al., 2023). The cells were also lysed to evaluate intracellular HBV DNA level by real time PCR.

**HBV spread assay**

PHH (PXB) cells were inoculated with HBV genotype C (PhoenixBio) at a very low inoculum (5 GEq/cell) in the presence of 4% PEG8000 for 16 h, followed by washing to remove the unbound viral particles, the cells were cultured for 42 days to allow viral spread. The culture supernatant was harvested every 6 days to quantify HBV DNA over time (Tsukuda et al., 2017).

**Indirect Immunofluorescence (IF) assay**

Immunofluorescence analysis was performed as previously describe (Ibrahim et al., 2021). Briefly the cells were fixed with 4% paraformaldehyde, permeabilized with 0.3% Triton X-100, incubated with anti-HBc (Invitrogen, PA5-16368), and then incubated with Alexa Flour555-conjugated secondary antibody, together with DAPI. Microscopic examination of HBcAg signal in HBV infected cells was conducted using fluorescence microscope.

**Immunoblot assay**

Protein detection through immunoblotting analysis was performed as previously described (Aly et al., 2016). The following primary antibodies have been used; mouse monoclonal E-cadherin antibody (Santa-Cruz, sc-8426), anti-β-actin (Sigma-Aldrich, A5441), anti-FOXM1 (Santa Cruz, sc-271746); and rabbit polyclonal anti-KIF4A (Invitrogen, PA5-30492), anti-NTCP (Sigma, HPA042727). For detection of NTCP level, the samples were incubated with Peptide-N-Glycosidase F (PNGase F) at 250 U to remove N-linked oligosaccharides from glycosylated NTCP before loading to SDS-PAGE (Gad et al., 2022).

**Extraction of surface proteins**

Cell fractionation using surface biotinylation assay to isolate the cell surface proteins was performed as previously described (Gad et al., 2022). The isolated surface fraction was then subjected to immunoblot assay as described earlier. E-cadherin (CDH-1) was used as internal control for the surface fraction (Gad et al., 2022).

**NTCP bile transporter assay**

The bile transporter activity into PHH (PXB) cells has been evaluated as previously described (Gad et al., 2022).

**DNA and RNA extraction**

Extraction of Intracellular HBV DNA and HBV cccDNA from the cells was performed as previously described (Ibrahim et al., 2021). Recovery of extracellular HBV DNA from the culture supernatant was conducted using SideStep Lysis and Stabilization Buffer (Agilent Technologies, 400900), while RNA extraction using the NucleoSpin® RNA XS Kit (MACHEREY-NAGEL) was performed according to the manufacturer’s guidelines.

**Quantification of viral RNA, HBV total DNA and cccDNA**

Following RNA extraction, RNA was reverse transcribed to cDNA as previously described (Gad et al., 2022). HBV pgRNA was quantified using Power SYBR green PCR master mix (Applied Biosystems) and normalized relative to GAPDH expression level (Ibrahim et al., 2021). RT-qPCR detection of HDV RNA and qPCR quantification of HBV (genotype D) DNA and cccDNA were primarily conducted as previously reported described (Gad et al., 2022). qPCR quantification of HBV DNA (genotype C) was conducted using the following primer-probe set (Liu et al., 2007); 5′-ACTCACCAACCTCTTGTCCT-3′, 5′-GACAAACGGGCAACATACCT -3′, 5′-FAM-TATCGTTGGATGTGTCTGCGGCGT-TAMRA-3′.

**Southern blot analysis**

Total intracellular HBV DNAs were detected using Southern blot technique as previously described (Ibrahim et al., 2021).

**ELISA quantification of secreted HBsAg and HBeAg**

Culture supernatants were harvested for detection of HBsAg as described previously (Gad et al., 2022), while ELISA evaluation of HBeAg was performed using Wantai commercial HBeAg ELISA kit (China, WB-2496) according to the manufacturer’s instructions (Yan et al., 2012).

**Assessment of cell viability**

The effect of the chemical compounds on cell viability was assessed using the Cell Proliferation Kit I (MTT, Roche), according to the manufacturer’s instructions.

**Data processing and statistical analysis**

The experiments were performed in triplicates or quadruplicates, and the means of data from 2-3 independent experiments were calculated and presented ± standard deviation (SD). Statistical significance was determined using Two-tailed unpaired student’s *t* tests (**, *P* < 0.01; ***, *P* < 0.001; NS, not significant).
