## Supplementary material for "RXR agonist S169 inhibits HBV/HDV entry *in vitro* by disrupting KIF4-dependent NTCP trafficking": Fig. S1

**Supplementary Figure Legend**

**Figure S1. Chemical structure of Oxaprozin and its RXR-modulating derivatives.**

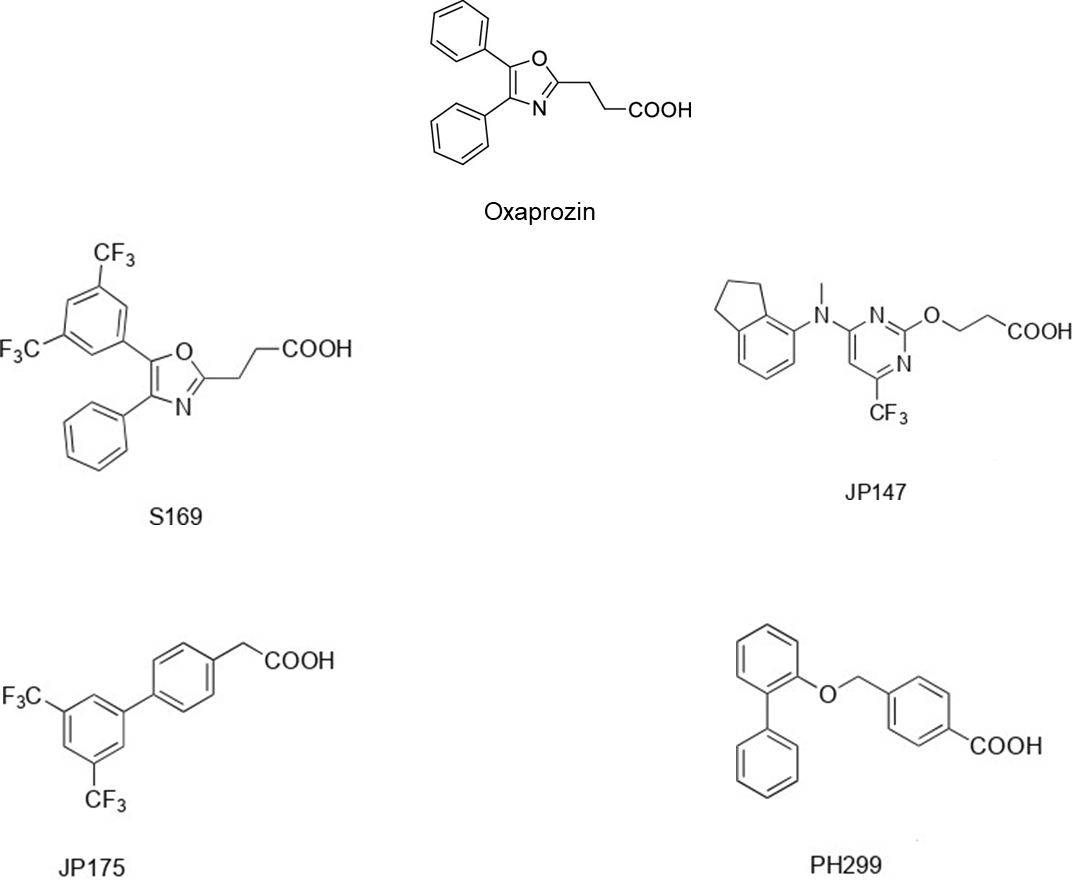
